# Pheromone receptor of the globally invasive quarantine pest of the palm tree, the red palm weevil (*Rhynchophorus ferrugineus*)

**DOI:** 10.1101/2020.07.31.230326

**Authors:** Binu Antony, Jibin Johny, Nicolas Montagné, Emmanuelle Jacquin-Joly, Rémi Capoduro, Khasim Cali, Krishna Persaud, Mohammed Ali Al-Saleh, Arnab Pain

## Abstract

Palm trees are of immense economic, sociocultural, touristic and patrimonial significance all over the world, and date palm-related knowledge, traditions and practices are now included in UNESCO’s list of the Intangible Cultural Heritage of Humanity. Of all the pests that infest these trees, the red palm weevil (RPW), *Rhynchophorus ferrugineus* (Olivier) is its primary enemy. The RPW is a category-1 quarantine insect pest that causes enormous economic losses in the cultivation of palm trees worldwide. The RPW synchronizes mass gathering on the palm tree for feeding and mating, regulated by a male-produced pheromone composed of two methyl-branched compounds, (4*RS*,5*RS*)-4-methylnonan-5-ol (ferrugineol) and 4(*RS*)-methylnonan-5-one (ferrugineone). Despite the importance of odorant detection in long-range orientation towards palm trees, palm colonization and mating, nothing regarding the molecular mechanisms of pheromone detection in this species is known. In this study, we report the identification and characterization of the first RPW pheromone receptor, *RferOR1*. Using gene silencing and functional expression in *Drosophila* olfactory receptor neurons, we demonstrate that *RferOR1* is tuned to both ferrugineol and ferrugineone and binds five other structurally related molecules. We reveal the lifetime expression of *RferOR1*, which correlates with adult mating success irrespective of age, a factor that could explain the wide distribution and spread of this pest. As palm weevils are challenging to control based on conventional methods, elucidation of the mechanisms of pheromone detection opens new routes for mating disruption and the early detection of this pest *via* the development of pheromone receptor-based biosensors.

## Introduction

The red palm weevil (RPW), *Rhynchophorus ferrugineus*, is the most damaging insect pest of palm trees and responsible for massive economic losses worldwide. Originating from South Asia, this invasive weevil has spread through the Middle East, Africa and the whole Mediterranean area since 1980. Its control relies mainly on the use of systemic insecticides, but ecofriendly management techniques, such as the inclusion of pheromone traps in integrated pest management strategies, have emerged. Pheromone traps are also used to monitor populations and new potential invasions, as this species poses a global threat.

The RPW pheromone is composed of two methyl-branched short-chain hydrocarbons, (4*RS*,5*RS*)-4-methylnonan-5-ol (ferrugineol) and (4*RS*,5*RS*)-4-methylnonan-5-one (ferrugineone), at a 9:1 ratio. It is produced by males and leads to coordinated mass attacks that often cause the collapse and death of the palm tree ^1^. Despite the fundamental importance of this pheromone in RPW mass attack and sexual reproduction as well as its use for insect control, nothing regarding the mechanisms of its recognition by these insects is known. It is hypothesized that this airborne pheromone is detected by a specific subclass of odorant receptors (ORs) specialized in pheromone detection. Indeed, most odorants are detected in insects *via* the activation of ORs, which act as gate-keepers of selectivity and sensitivity^2,3^. These ORs are heptahelical transmembrane proteins embedded in the dendritic membrane of olfactory receptor neurons (ORNs) housed in olfactory sensilla on the antennae. Compared to G-protein-coupled receptors (GPCRs), insect ORs present an inverse topology, Ni_in_–C_out_, and insect ORs function as ligand-gated nonselective cation channels^4,5^ *via* heteromeric complexes formed by an odorant-specific OR protein and a highly conserved coreceptor (Orco) ^6–10^. Electrically encoded messages are conveyed to the central nervous system in the antennal lobe, where the neural signal is decoded^10,11^. Thus, insect ORs are the main gateway to sensing the world of volatile chemicals ^2,10,12^. Although many OR sequences from a range of insect species, including that of the RPW ^13^, are now available, information about the functional role of ORs remains limited to the OR repertoires of a few model organisms, such as the fruit fly *Drosophila melanogaster* ^14^, and ORs tuned to particular signals, such as moth sex pheromones^3,11,15,16^. Notably, the response spectra of only a handful of ORs have been studied in beetles (Coleoptera)^17,18^, even though they constitute the most species-rich insect order. In particular, no RPW ORs have been functionally characterized. In this study, we specifically screened for RPW aggregation pheromone receptors by selectively silencing ORs highly expressed in the antennae using RNA interference and assessing changes in pheromone detection using electrophysiological recordings. By doing so, we identified one OR, *RferOR1*, as the best candidate pheromone receptor. We then heterologously expressed it in *Drosophila* ORNs and established its response spectrum to a large panel of weevil and palm beetle pheromone components. *RferOR1* was best activated by ferrugineol and ferrugineone and slightly less activated by five other structurally related molecules, thus demonstrating its functional role as an *R. ferrugineus* aggregation pheromone receptor. Subsequently, we modeled the 3-dimensional structure of *RferOR1* and proposed binding sites.

This work represents an essential step in our understanding of RPW biology. This study identifies an OR as a new target for the development of pheromone receptor-based biosensors for the early detection of RPWs in the field.

## Results

### RferOR phylogeny and expression analyses

Thanks to sequencing of the RPW antennal transcriptome, we previously annotated 71 non-redundant candidate *R. ferrugineus* ORs ^13^. A maximum likelihood (ML) tree was constructed with Curculionidae and Cerambycidae OR sequences and revealed that the 71 *R. ferrugineus* ORs were clustered in four (1, 2, 5 and 7) of the seven major OR groups already described ^18^, with the largest number of RferORs found in group 7 (35 ORs), followed by groups 1 (14 ORs), 2B (10 ORs), 2A (8 ORs) and 5A (4 ORs), which is slightly dissimilar to the OR distribution in the other Curculionidae species used in the analysis (Fig. 1). *R. ferrugineus* lacked ORs in groups 3, 4, 5A and 6. Gene expansions and species-specific clusters were identified in the tree (Fig. 1). The phylogenetic analysis also indicated that several *R. ferrugineus* ORs display a 1:1 orthologous relationship with *Sitophilus oryzae* ORs, with >95% bootstrap support (Fig. 1). The functionally characterized pheromone receptors (PRs) from *I. typographus* ^18^ and *M. caryae* ^17^ were located in different clades (Fig. 1). Whereas the two ItypPRs were found in clade 7, two of the three *McarPRs* were found in clade 2B, and the third (*MacrOR3*) was in clade 1, clustered with *RferOR41* (82% bootstrap support). Using RT-PCR, we next verified the expression of each of the *RferOR* genes in the antennae of females and males and in other tissues in males (Fig. S1). We identified seven *RferOR* genes ubiquitously expressed in all tissues studied, five with no expression in the wings but expression in all other body parts, twenty with antennae-specific expression, one candidate male-biased OR (*RferOR32*) and the remaining *RferOR* genes with various expression patterns (Fig. S1). Then, we quantified the expression levels of the 20 antennae-specific *RferORs* (hereafter named *RferOR1* to *OR20, Dataset S1*) in male and female antennae. They displayed various expression levels in the antennae of laboratory-reared animals (Fig. S1), with relative expression levels ranging from 0.001 to 5.24 (normalized to *tubulin* and *β-actin* gene expression) (Table S2). A group of eight *RferORs* (*RferOR1* to 8) exhibited visibly higher expression levels than the others (Fig. S1). Five of them belong to clade 7, two belong to clade 2B and one belongs to clade 1 (Fig. 1).

**Fig. 1.**
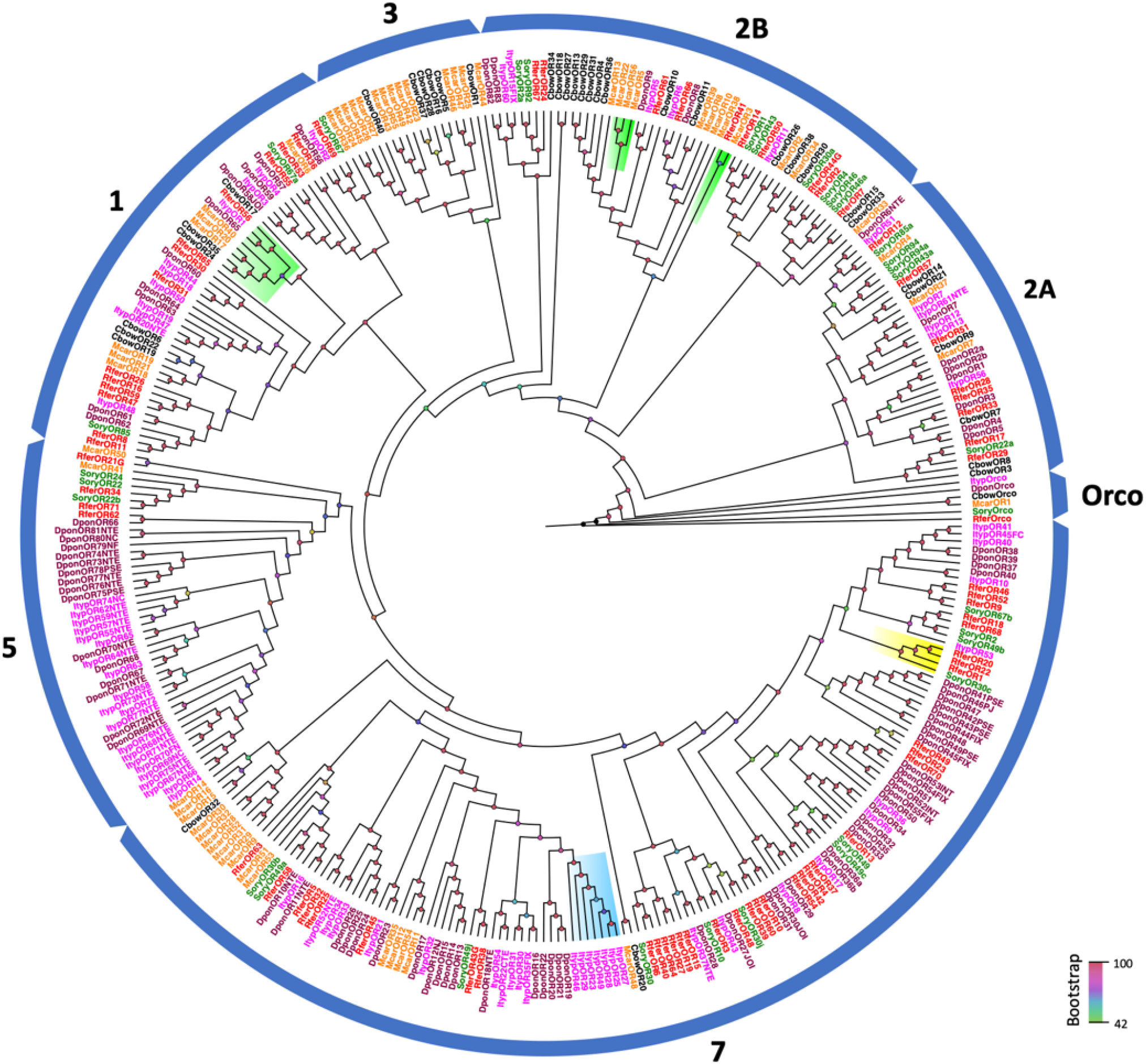
Maximum likelihood consensus tree of ORs from Coleoptera. The tree included OR amino acid sequences of *R. ferrugineus* (red) and other coleopterans retrieved from GenBank including *Colaphellus bowringi*, Cbow (black); *Ips typographus*, Ityp (magenta); *Dendroctonus ponderosae*, Dpon (maroon); *Megacyllene caryae*, Mcar (orange) and *Sitophilus oryzae*, Sory (Green). The branch containing Orco was used as an outgroup. Based on the phylogeny, representative coleopteran OR clades are indicated with blue arcs and numbers representing the clades. The functionally characterized ORs are highlighted; *R. ferrugineus* (yellow), *Ips typographus* (blue) and *Megacyllene caryae* (green). Bootstrap values are indicated at the nodes with a color gradient from green (42 %) to red (100 %).

### *RferOR1* is overexpressed in response to pheromone pre-exposure

To identify the RferORs responsible for detection of the aggregation pheromone, we next measured whether the expression levels of any RferORs were regulated by pre-exposure to a synthetic pheromone blend. Overall, changes in the expression patterns of all 20 ORs in the pheromone pre-exposed group compared to the laboratory sample control group were relatively minor (Table S2). Among the twenty ORs analyzed, most ORs showed no change in expression or lower expression compared to that in non-exposed laboratory RPW controls (Table S2), whereas *RferOR1* was found to be marginally upregulated in pheromone pre-exposed RPWs (Fig. 2a). Interestingly, the expression level of *RferOR1* upon pre-exposure in laboratory RPWs was on par with that in the field-collected RPWs (Fig. 2b), which are assumed to be under constant pheromone exposure in the field. *RferOR1* expression in the pheromone-exposed and field RPW groups was higher than the expression of all other 19 RferORs (*RferOR2* to *RferOR20*), based on Waller-Duncan analysis (Fig. 2a).

**Fig. 2.**
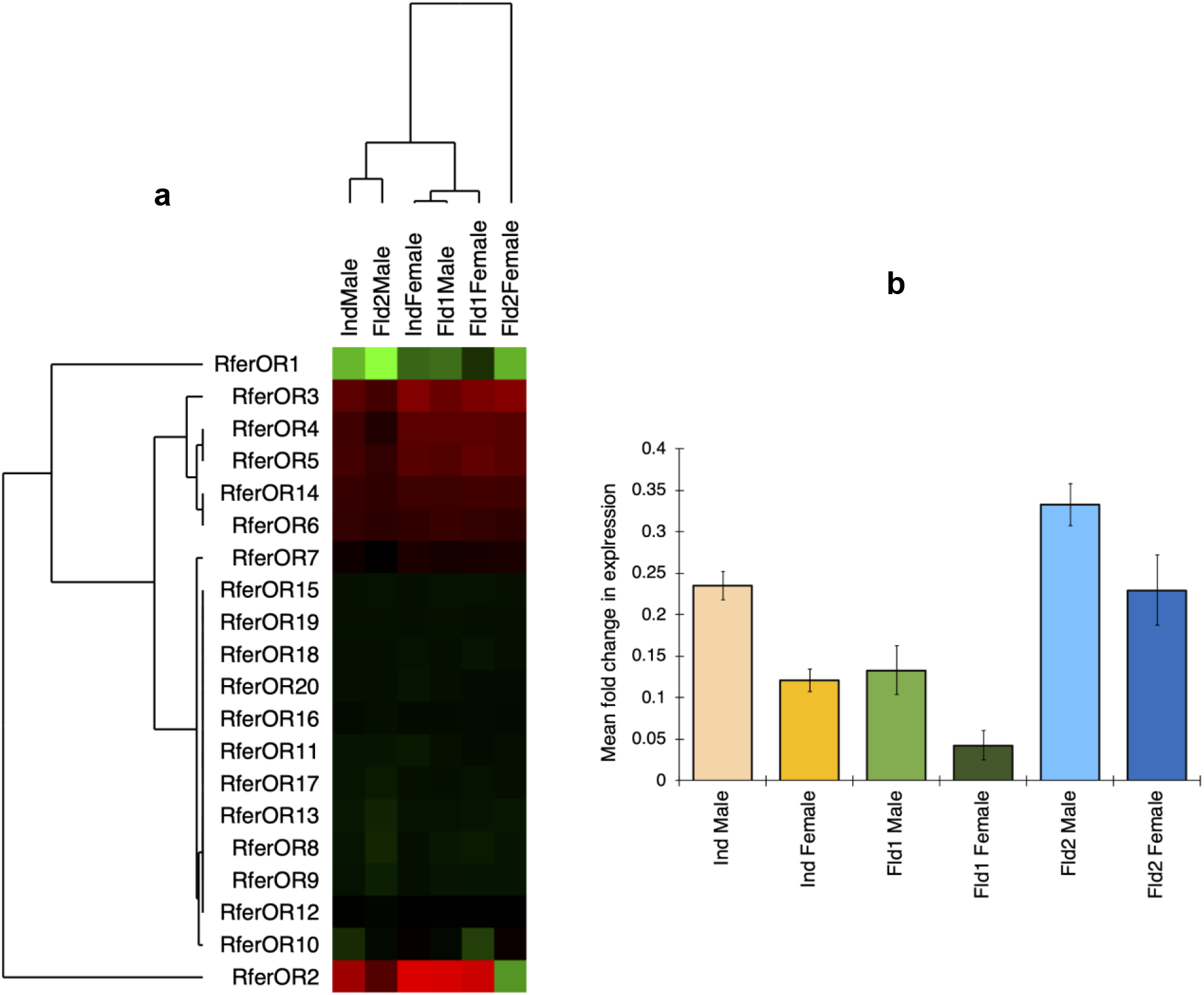
*RferOR1* is overexpressed in response to pheromone pre-exposure. **(a)**. Expression analysis of *R. ferrugineus* antennal specific ORs using RT-qPCR, with and without pheromone pre-exposure. The hierarchical cluster analysis shows the fold changes in the expression pattern of 20 antennae specific ORs with respect to 20-days old male and female lab insects (*Supplementary Methods*). The different test groups insects pre-exposed with commercial aggregation pheromone (abbreviated as *Ind*) and samples collected from two different date palm fields in Saudi Arabia (*Fld1* and *Fld2*). The data represents log-transformed RT-qPCR 2^-Δ△C_T_^ values. The heatmap colors represent expression level form highest (green) to lowest (red) expression. (**b**). Mean fold changes in *RferOR1* expression in pheromone pre-exposed and field-collected RPWs (*Supplementary Methods*). The relative expression of all other antennal specific ORs provided in Table S2.

### *RferOR1*-silenced RPW adults exhibited a reduced response to pheromone

Based on the results of *RferOR* expression analyses, we focused on the six most highly expressed *RferORs* and further investigated their potential contribution to pheromone detection. To do so, we used gene silencing with RNAi. Injection of dsRNA successfully knocked down expression of the six *RferOR* genes, with knockdown efficiencies compared to non-injected controls of between 75 and 80% (Fig. 3a).

**Fig. 3.**
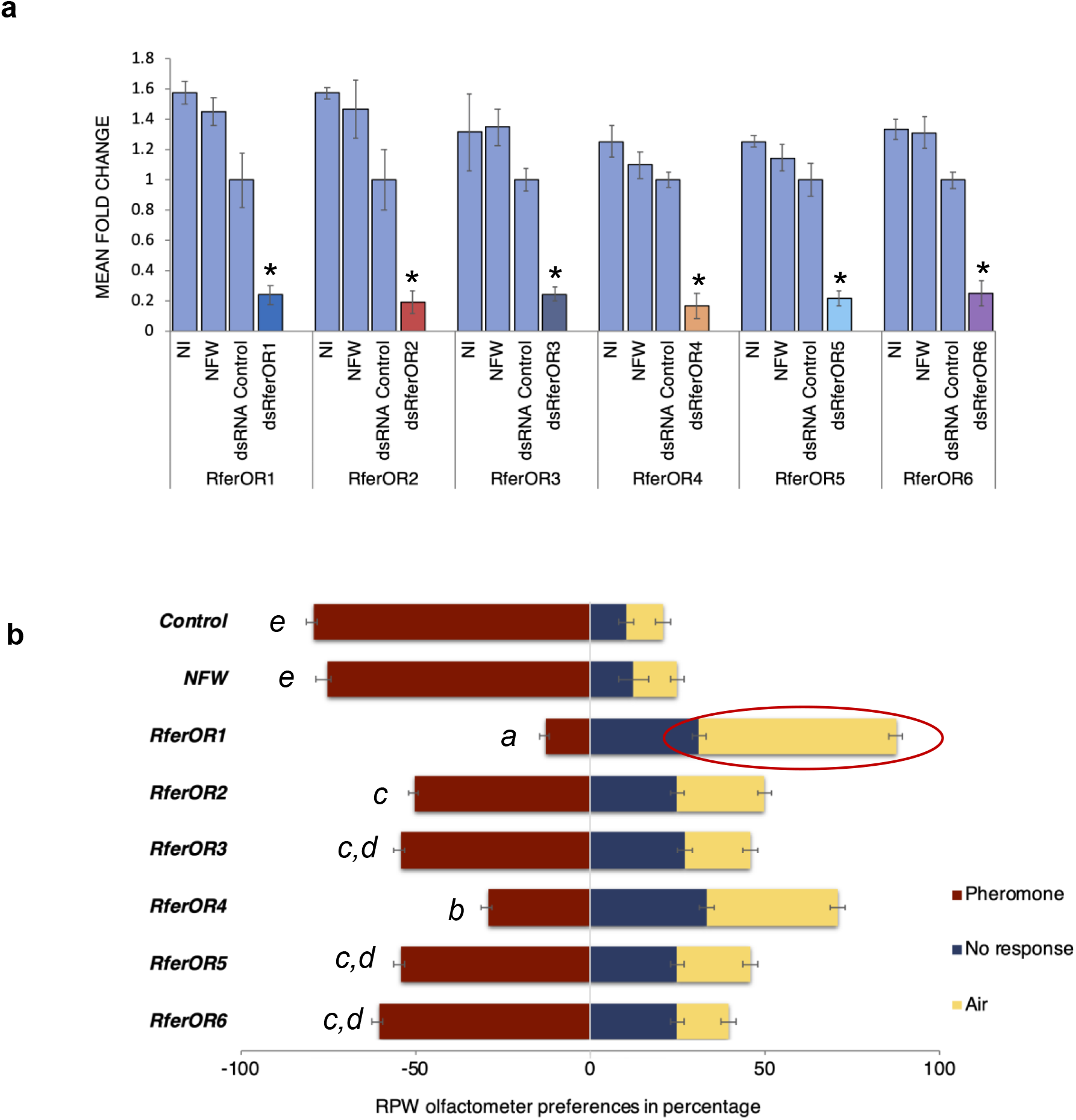
RNAi experiment and behavioral assays. **(a)**. Effects of RNAi based silencing on *RferOR* expression, measure using qRT-PCR. Data presented as mean fold change in each *RferOR* expression (2^-ΔΔCt^ values) normalized with dsRNA control (abbreviations, NI: non-injected, NFW: Nuclease free water injected, dsRNA Control: negative control dsRNA injected and dsRferOR: respective RferOR dsRNA-injected). (*) represents the statistical significance measured at *P* < 0.05 and error bars represents SEM (*P* < 0.05; one-way ANOVA with LSD) (see Table S3). (**b**). Effects of RNAi on behavior. Olfactometer preferences exhibited by OR-silenced (RferOR dsRNA-injected) RPWs against NI (not-injected) and NFW-injected insects to pheromone (*SupplementaryMethods*). Responses were provided as ‘towards pheromone’ (red), ‘no response’ (blue), and ‘towards air’ (yellow) expressed as percentage of the total (*n* = 16). *RferOR1* silenced adults show significant response ‘towards air’ is highlighted with an oval circle. Error bars represent SEM and letters (*a, b, c, d*, and *e*) represent five homogenous subsets identified by Waller-Duncan statistical analysis.

In the behavioral assays, *RferOR1*-silenced animals showed significant behavioral changes in response to a commercial aggregation pheromone compared to the behavior of controls. We found that 56.25% of *RferOR1-silenced* RPWs were unable to respond to pheromone stimuli, as the adults moved away from the pheromone source towards clean air, and 31.25% had no response at all (Fig. 3b). Only 12.5% of the silenced animals were attracted to the pheromone, a significantly smaller proportion than that of the non-injected controls (79.17%) and NFW-injected controls (75%) (*P* <0.001; Table S3). *RferOR1*-silenced insects also had significantly different responses than other *RferOR*-silenced insects (*P*=0.001). Waller-Duncan analysis for homogenous subsets confirmed *RferOR1* and *RferOR4* as separate subsets *a* and *b*; *RferOR2, RferOR3, RferOR5*, and *RferOR6* as subsets *c* and *d*; and the controls NI and NFW as subset *e* in response to commercial aggregation pheromone (Fig. 3b). Similarly, with their reduced response to pheromone, *RferOR1-*silenced insects were grouped as a homogenous subset. Results of the olfactometer study results revealed that the most altered pheromone behavioral response was obtained when *RferOR1* was silenced.

The EAG results showed that *RferOR1*-silenced insects had a significantly reduced response to ferrugineol (Fig. 4a, b), with an amplitude of 2.57 (± 0.67) mV compared to that of 8.12 (± 0.42) mV in control insects (*P* < 0.001; Table S4). The mean response of *RferOR1-silenced* insects to ferrugineone was also slightly reduced but not significantly different from that of control animals. All other *RferOR*-silenced experimental groups showed unaltered EAG responses to ferrugineol, ferrugineone and the host volatile ethyl acetate, with no significant differences compared to control insects (Fig. 4a, b).

**Fig. 4.**
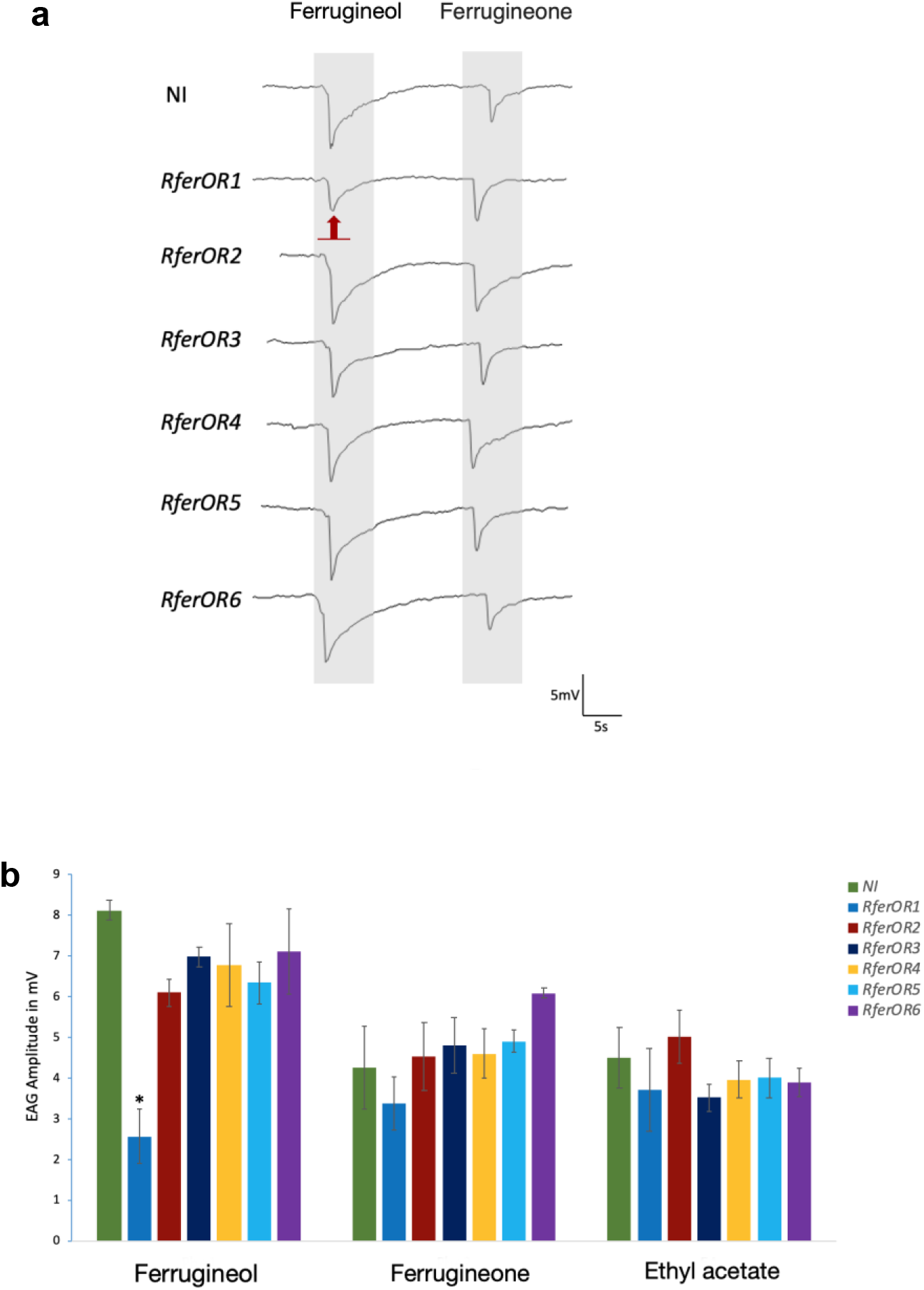
*RferOR1*-silenced RPW adults exhibited a reduced response to pheromone. **(a**). Representative EAG recordings from the *RferOR*-silenced insects compared with non-injected (NI) weevils. Grey columns represent stimulus duration. The red arrow indicates a reduction in EAG response in *RferOR1*-silenced insects when stimulated by ferrugineol. (**b**). EAG amplitudes measured from RNAi *RferOR*-silenced and non-injected (NI) insects when stimulated with 200 ng on the filter paper of the aggregation pheromone components ferrugineol and ferrugineone and with volatile ethyl acetate (mean values ± SEM).* *P*<0.001 (one-way ANOVA followed by a Tukey’s HSD method) indicates a significant reduction in *RferOR1*-silenced insect’s response to ferrugineol compared to NI (see Table S4). Error bars represent SEM.

### Transgenic *Drosophila* olfactory neurons expressing *RferOR1* responded to ferrugineol and ferrugineone

Based on its high expression level and marginal overexpression upon pheromone pre-exposure and the reduced antennal and behavioral responses to pheromone following its knockdown by RNA interference, *RferOR1* appeared to be the best candidate *R. ferrugineus* pheromone receptor. For further confirmation, we heterologously expressed an *RferOR1* transgene in *D. melanogaster* ORNs from the at1 sensilla deprived of endogenous OR67d, which is tuned to cVA ^19^. We first verified the lack of response to cVA in transformed *Drosophila* at1 ORNs, confirming the absence of OR67d (Fig. 5a). Then, the ORNs were stimulated with high doses of ferrugineol, ferrugineone and a range of pheromone compounds from other species of weevils or palm beetles (Table S5) and structurally related compounds. We found strong responses to ferrugineol and ferrugineone (mean responses of 106 and 95 spikes.s^-1^, respectively) and minor responses to nonan-5-ol, nonan-5-one, oct-1-en-3-ol, (*E*)-oct-2-en-4-ol and 5-methyloctan-4-one (Fig. 5a). We next carried out dose-response experiments for the four most active compounds. Ferrugineol appeared to be the best RferOR1 agonist, although ferrugineone starting at a dose of 100 ng also induced significant responses. Nonan-5-ol and 5-methyloctan-4-one induced responses, but only at doses of 1 and 10 μg (Fig. 5b, c). With those experiments, we thus confirmed our hypothesis that *RferOR1* is involved in pheromone detection in *R. ferrugineus*.

**Fig. 5.**
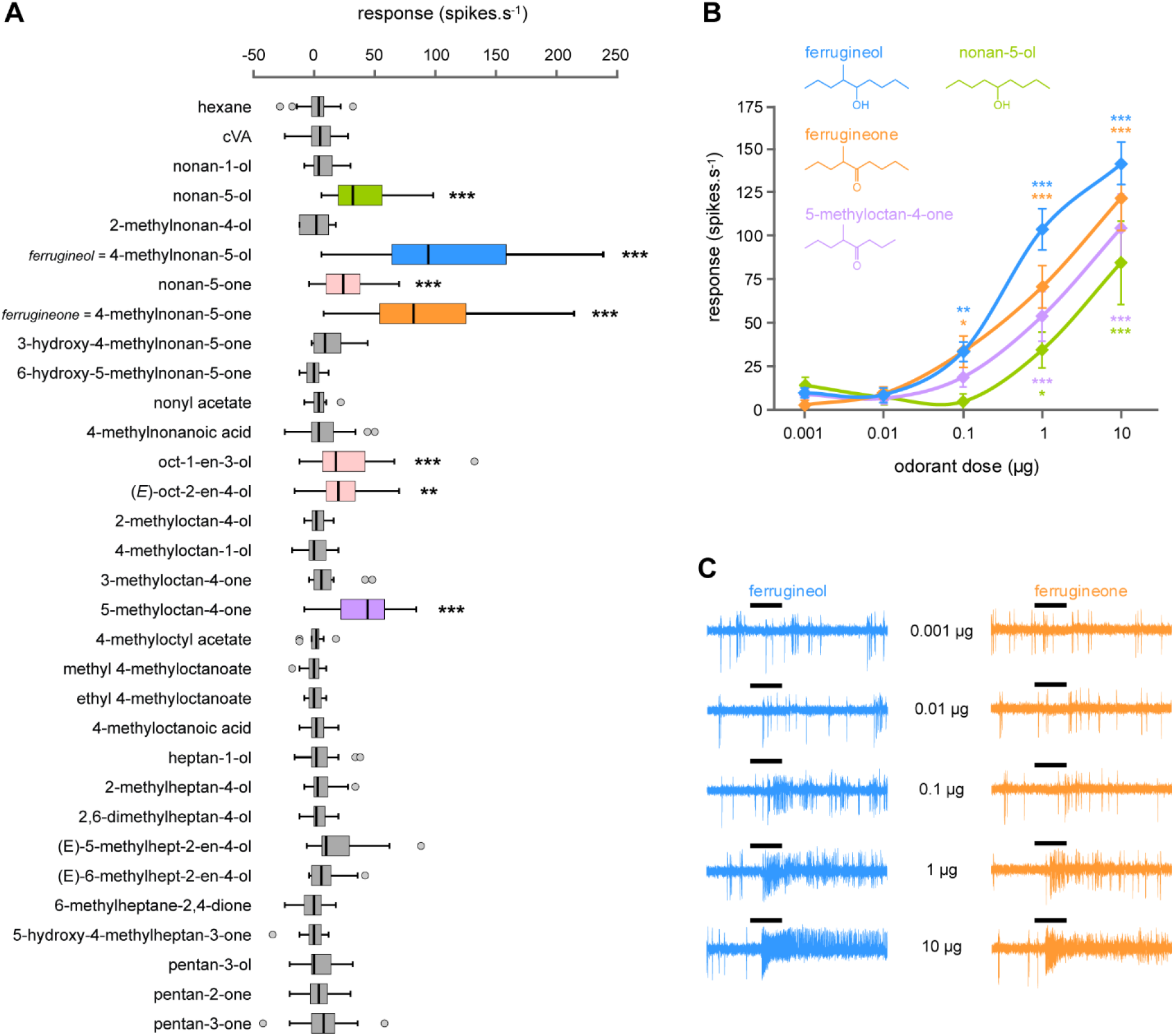
RferOR1 is activated by ferrugineol, ferrugineone and five other structurally related compounds. (**a**) Action potential frequency of *Drosophila* OSNs expressing RferOR1 when stimulated with a panel of pheromone compounds and related chemicals (10 μg loaded in the stimulus cartridge). Box plots show the median and the first and third quartiles of the distribution (*n* = 13-49). ** *P*<0.01, *** *P*<0.001, significantly different from the response to solvent alone (Kruskal-Wallis ANOVA followed by a Dunn’s post-hoc test). (**b**) Dose-response curves of *Drosophila* OSNs expressing *RferOR1* to four active compounds. Data represented are mean action potential frequencies ± SEM (*n* = 9-15). * *P*<0.05, ** *P*<0.01, *** *P*<0.001, significantly different from the response to solvent alone (Kruskal-Wallis ANOVA followed by a Dunn’s post-hoc test). (**c**) Single sensillum recordings obtained for a *Drosophila* at1 OSN stimulated with increasing doses of ferrugineol and ferrugineone. Black bars represent the duration of the stimulus (500 ms).

### Structural modeling and molecular docking of RferOR1

The 1233 bp open reading frame (ORF) of *RferOR1* encodes a protein of 411 amino acid residues whose structure was modeled (Fig. 6a). We identified a total of ninety-four pockets, with the main binding pocket (the probable active site) predicted to be made up of 28 amino acid residues (Table S6), of which fourteen (Table S7) are hydrophobic (50%), seven are hydrophilic (25%), two are positively charged (7.14%) and five are negatively charged (17.86%) (Fig. S2). In RferOR1, the channel to the active site had a distinct entry mouth and exit mouth. The pocket had two mouth openings (Table S6); the residues making up the entry mouth (Table S7) were D77, K78, D298, K301, V302, and D308, while those making up the exit mouth were N174 and S176 (Table S6 and Fig. S3 a, b). These features of the active site are summarized in Table S7.

**Fig. 6.**
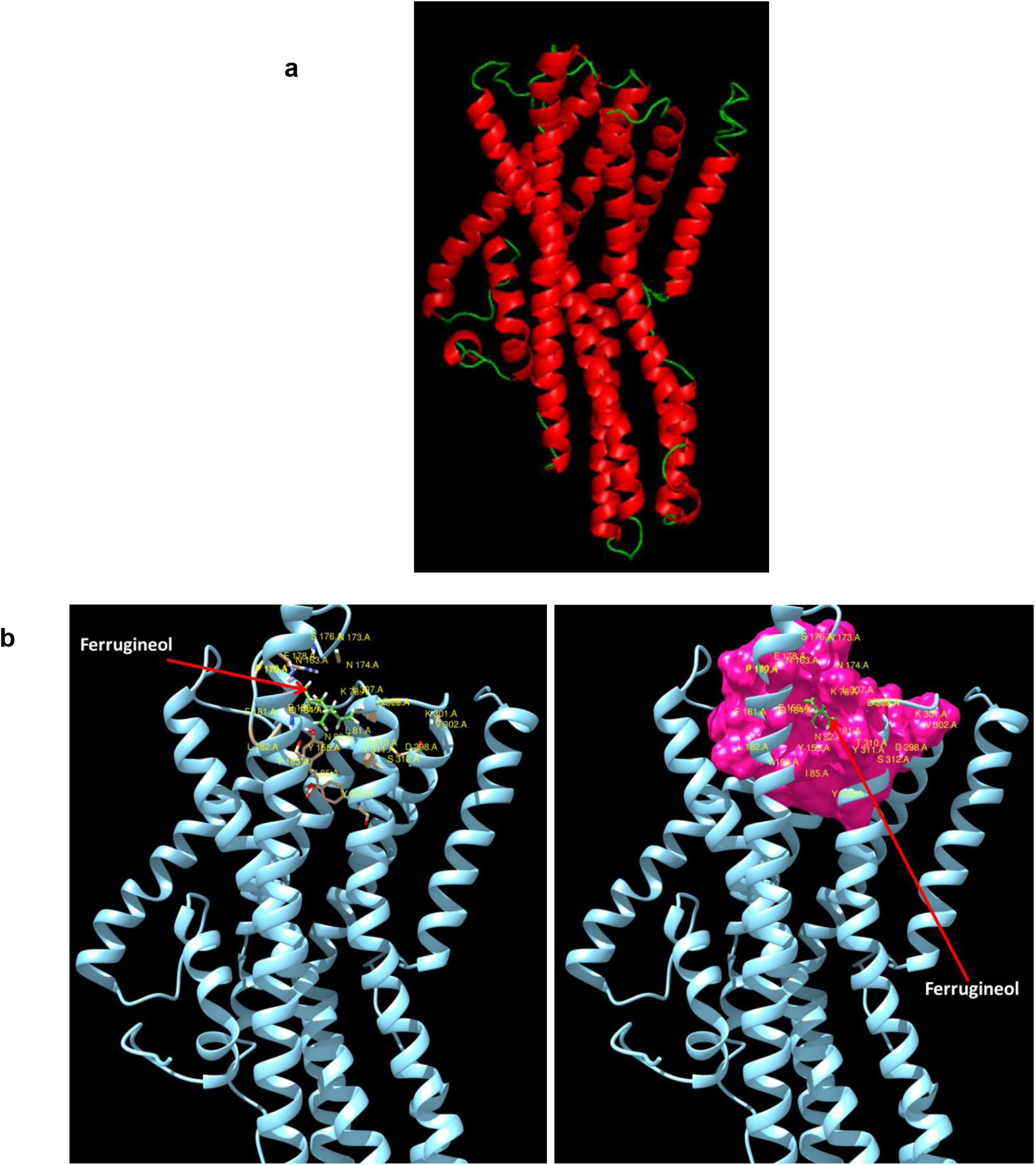
Structural modeling and docking of red palm weevil pheromone receptor, RferOR1. **(a**). Cartoon presentation - *Helix and Loop* of the structure. The figures were rendered using molecular visualization software PyMOL (Schrodinger). **(b**). Binding of aggregation pheromone 4-methyl-5-nonanol (ferrugineol-labelled in green) to the binding pocket of RferOR1 (left and right). In right figure, the pocket is shown in hydrophobicity surface representation and colored in dark pink.

We then conducted docking studies using the different compounds used to challenge RferOR1 expressed in *Drosophila* (Table S8), five additional related compounds and VUAA1, an Orco agonist ^20^. After these molecules were docked against the protein-binding site, the data were analyzed, as described in the Methods section. For each ligand, the most energetic binding pocket-binding mode was selected, and the binding energy (Table S9) was used to define the protein binding affinity towards a given molecule.

With the exception of VUAA1, all other molecules bound within the active site of the protein, giving negative binding energies (Fig. 6b). The lower the binding energy was, the stronger the binding was (higher affinity) and *vice versa*. As an example, Figure 6b illustrates the binding of a ferrugineol molecule to the active site of the receptor. All tested ligands except VUAA1 were able to fully penetrate the binding pocket (Fig. S4). Comparing the two aggregation pheromone components, the receptor showed slightly higher affinity towards ferrugineol (binding energy, −31.73 kcal/mol) than ferrugineone (−28.89 kcal/mol) did. Within the active site, seven (out of the 28) residues made a direct interaction with ferrugineol; these residues were L81, Y155, N174, E178, P180, Q184, and Y311 (Table S10 and Fig. S5). N174 was one of two residues (the other one being S176) that make up the active site exit mouth (Table S6 and Fig. S5).

### *R. ferrugineus* pheromone receptor expression was observed throughout the life cycle

We observed a slight difference in *RferOR1* expression patterns between male and female antennae; however, the values were not significantly different and thus did not show sex-biased expression, in accordance with our tissue-specific expression analysis (Fig. 7a). Importantly, *RferOR1* expression was observed throughout the life cycle (0-60 days), with maximal expression at 20 days. At 60 days, *RferOR1* expression was higher than that in newly emerged RPWs (0 days) (Fig. 7b). We recorded the adult RPW response to the aggregation pheromone using EAG data collected from RPWs at different ages, which showed a significantly higher response to ferrugineol and ferrugineone in both 20-day-old males and females than in young and old adults (*P*<0.01). The EAG results showed that 60-day-old male and female adults responded to the aggregation pheromone with an amplitude similar to that of newly emerged RPWs (0 days) (*P* = 0.072 in males and *P* = 0.409 in females for ferrugineol, *P* = 0.362 in males and *P* = 0.668 in females for ferrugineone) (Fig. 7c).

**Fig. 7.**
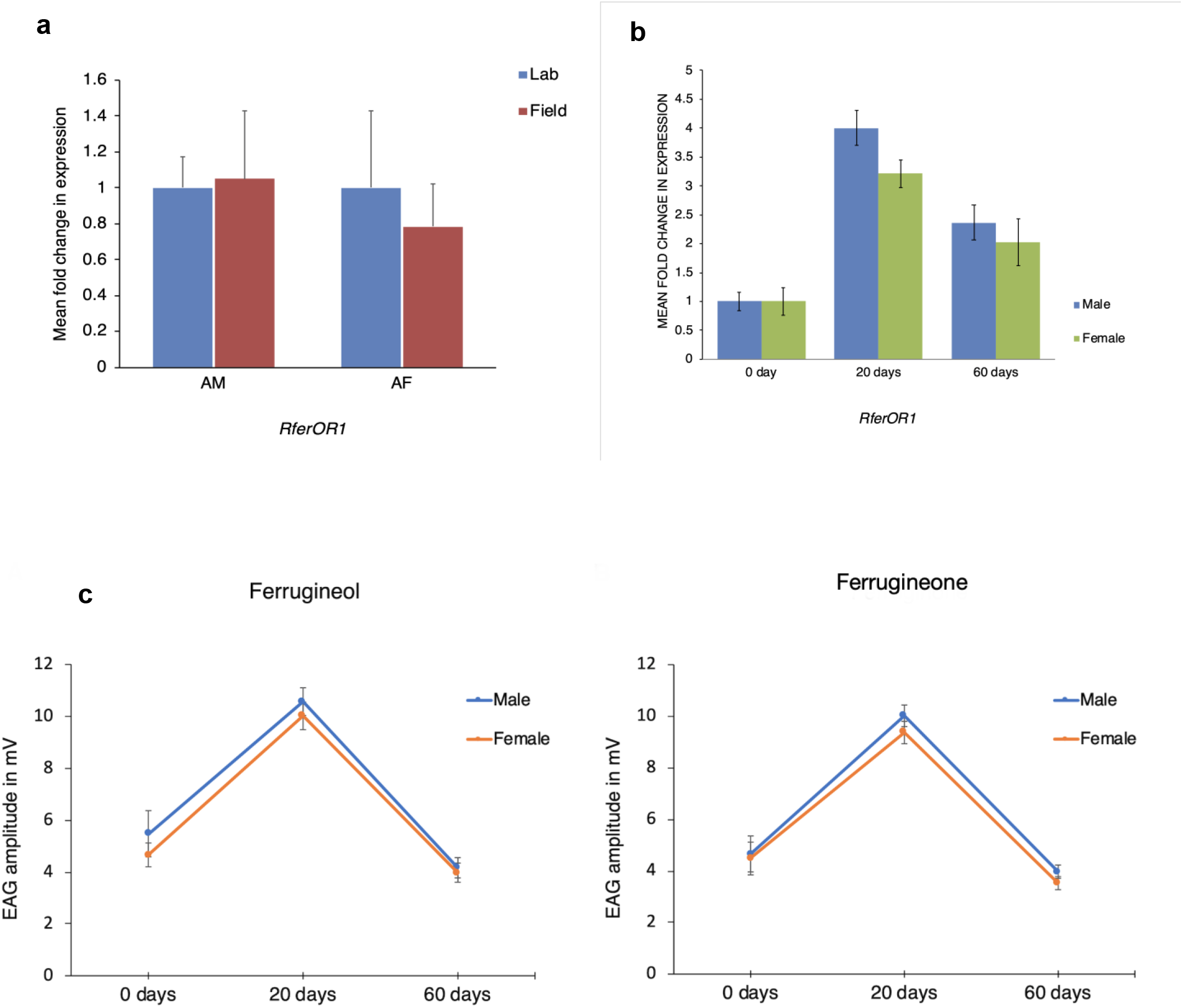
*RferOR1* expression and antennal response to the pheromone according to age. (**a**). Relative expression of *RferOR1* in antennae from RPW male and female adults (RT-qPCR) (fold changes compared to the expression level in lab-reared male antennae). (**b**). *RferOR1* relative expression in antennae of 0, 20, and 60-day-old RPW male and female adults (fold changes compared to the expression level in antennae of 0-day old lab-reared males). **c.** EAG recordings (*mV*) in response to ferrugineol and ferrugineone in 0 days, 20 days, and 60 days RPW adults. The error bars represent SEM.

## Discussion

In recent years in the Anthropocene epoch, many unpredicted and irreversible changes in insect pest distribution and behavior have occurred, and some insect species have, with human help, spread rapidly across the world. The phenomenal expansion and global spread of the red palm weevil in almost all dominant palm tree-growing countries over the last three decades has recently resulted in its attainment of category-1 pest status ^21^ as a consequence of the commercial exchange of palm trees worldwide ^22–24^. In the palm weevil, both long-range attraction and host colonization for feeding and mating are regulated by an aggregation pheromone^24,25^. Understanding the molecular mechanism of pheromone reception and elucidating the functional role of pheromone-specific ORs and odorant-binding proteins (OBPs) may help to design novel pest management solutions^26,27^. In this study, from 71 candidate RPW ORs, we identified one OR, *RferOR1*, as a pheromone receptor in *R. ferrugineus*. We demonstrated pheromone response disruption through RNAi-based gene silencing and confirmed the functional role of *RferOR1* in pheromone detection through heterologous expression in *Drosophila*. These findings provide the first insights into the molecular basis of chemoreception in a palm weevil species. They also represent a significant advance towards a complete understanding of olfaction in Coleoptera. While insects from this order, including phytophagous beetles of economic importance, pathogen vectors, predators and beneficial insects with key roles in the cycle of organic matter, exhibit a wide range of ecologies, it is surprising that only five other ORs in this order have been functionally characterized to date^18,28^.

Because of the paucity of data on Coleoptera ORs ^28^, we had to rely on several approaches to select ORs possibly involved in pheromone detection. First, we employed a phylogenetic approach to identify a coleopteran-specific pheromone receptor (PR) clade. However, the five Coleoptera ORs described as PRs did not cluster in the same clade in the phylogenetic tree (Fig. 1), precluding the identification of good RPW PR candidates. In fact, it has been proposed that coleopteran PRs are highly divergent and have acquired unique functions related to different pheromone compounds, contrary to lepidopteran PRs, which cluster in specific clades in accordance with structural similarity among pheromone compounds^28,29^. As PRs in insects are usually characterized by high transcript abundance and specific expression in the antenna, we followed these criteria to select candidate ORs *via* tissue expression studies and quantification of relative OR expression. Thereafter, we targeted the ORs that were highly and uniquely expressed in the antennae for further studies. We used another criterion, transcript upregulation upon exposure to pheromone stimuli. With all these criteria, we selected six highly expressed and pheromone-induced ORs for gene silencing. Perfect correlation between the reduction in *RferOR1* transcript level and the very modest antennal responses to the pheromone suggested that *RferOR1* is a pheromone receptor. We ultimately confirmed by heterologous expression in *Drosophila* ORNs that *RferOR1* is a receptor sensitive to ferrugineol and ferrugineone. In doing so, here, we provide the first evidence that *Drosophila* ORNs can be used efficiently for the study of ORs from Coleoptera, in addition to Diptera and Lepidoptera. Indeed, the function of previously characterized Coleoptera pheromone receptors was determined *via* their expression in *Xenopus* oocytes coupled to a two-electrode voltage clamp, a method that proved successful for ligand identification of ORs from species from a range of insect orders. Our work not only provides another tool for Coleoptera olfaction studies but also reveals that the ORN machinery, especially Orco, is functionally conserved across 440 MY of insect evolution ^30^.

Notably, at high pheromone doses (10 μg), *Drosophila* ORNs expressing *RferOR1* showed continuous spiking after stimulation. One explanation for this observation is that *Drosophila* at1 sensilla are devoid of adequate pheromone-degrading enzymes. Without pheromone degradation in these sensilla, the ORNs would be continuously stimulated. Similar effects of prolonged stimulation were observed in earlier studies using expression in *Drosophila*^31,32^. In addition, RferOR1, when expressed in *Drosophila* ORNs, responded to both components of the pheromone, whereas *RferOR1*-silenced RPWs presented an altered response to ferrugineol alone. A possible explanation is that *Drosophila* ORNs are not equipped with the correct OBPs for discrimination of the two compounds, as RPW sensilla are.

The specificity of RferOR1 towards the RPW pheromone was further assessed by challenging *Drosophila* ORNs with a wide range of compounds of various chain lengths (5C to 9C) and functional groups (Fig. 5). Fly ORNs presented a high sensitivity to the two components of the RPW aggregation pheromone (Fig. 5b) and sensitivity to a lesser extent to five structurally related compounds (nonan-5-ol, nonan-5-one, oct-1-en-3-ol, (*E*)-oct-2-en-4-ol and 5-methyloctan-4-one). One of these compounds, nonan-5-ol, is a component of the pheromone of *Metamasius hemipterus*, another Rhynchophorinae ^33^. RPW and *M. hemipterus* are not sympatric species, however, suggesting that nonan-5-ol does not interfere with RPW pheromone communication in the wild. Similarly, (*E*)-oct-2-en-4-ol is a pheromone component of the related species *R. palmarum*, whose native range extends from Central to South America. Of importance, none of the tested pheromones from weevil species living in sympatry with the RPW was detected by *RferOR1*, suggesting the maintenance of species isolation.

Interestingly, *RferOR1* was found to be expressed throughout the RPW lifespan, with its highest expression observed in adults at 20 days of life, correlating with the highest EAG signal, which was recorded at the same age. *RferOR1* expression was also observed in young and old (sixty-day-old) adults, which responded normally to the aggregation pheromone (Fig. 7b). This phenotype correlates well with the previous observation that adult mating success is independent of age ^34^. As pheromone reception and response are directly linked to reproductive success, this feature might have helped the phenomenal expansion of palm weevils worldwide.

Our docking studies based on binding energies defined the affinity of RferOR1 towards the RPW pheromone (Table S9). We decoded pheromone-RferOR1 interactions using *in silico* docking, and the results predicted that RferOR1 responds to ferrugineol and ferrugineone with high binding affinity, as observed experimentally in the *Drosophila* system. The difference in binding energies between the two aggregation pheromone compounds was not significant, but the binding energy towards ferrugineol was slightly higher than that towards ferrugineone (Table S9). This suggests that the active site features of RferOR1 that respond to both pheromone compounds are quite similar. It will be interesting and useful in the future to characterize the binding pockets to test the above hypothesis. Moreover, the critical amino acid residues involved in the formation of ferrugineol–RferOR1 complexes need to be validated by subsequent site-directed mutagenesis and fluorescence binding assays. Differences in affinities of the receptor to different ligands could also be attributed to the size and nature of the compounds tested with respect to the size and nature of the active site of RferOR1 (Table S9 and Figs. S2, S3). For example, a molecule such as ferrugineol (MW = 158.28) can fully access the active site of the receptor (Fig. S4) with a binding energy of −31.73 kcal/mol (Table S9). However, the molecule VUAA1 (MW = 367.47), which is 2.3-fold larger in size than ferrugineol, cannot access the pocket (Fig. S4), giving a positive binding energy of 30.16 kcal/mol (Table S8).

The primary purpose of our RferOR1 docking experiment was to obtain acceptable models of OR-ligand complexes (especially RferOR1-ferrugineol), for which no experimental structures are available, and to predict potential binding pockets and the active site of RferOR1. Because several studies indicated that subtle changes in the binding site compositions of ORs result in differential odorant binding and odor detection^30,35^, a lack of information about the deorphanized insect ORs in the database may lead to the poor prediction of protein affinity towards a given ligand ^35^. Hence, the results of docking screening based on binding energies should be validated and compared with the results of experimental functional studies ^36^. Our results, which suggest olfactory receptors of more insect species that need to be deorphanized, will be useful for increasing the accuracy of predictions with regards to specificity and selectivity across a wider repertoire of ORs, and might be comparable to the results of experimental functional expression studies. The data presented here characterizing the RferOR1 active site could be used in the future to investigate the roles of its constituent amino acid residues in ligand binding, but most importantly, these data could be used to design mutant variants that are more potent than the wild-type receptor for use as biosensors for early detection of the palm weevil in date palm fields. Preliminary energy analysis around the active site of the receptor using the Prediction of Protein Mutants Stability Changes (PoPMuSiC) server as described ^37^ indicated that mutation of twelve (out of the twenty-eight binding pocket residues) may produce stable mutants (Table S6). Looking back at the OR phylogenetic tree, we observed that *RferOR1* belongs to a monophyletic clade that groups other curculionid ORs, such as OR from *S. oryzae* (*SoryOR30c*) and two other antennae-enriched RPW ORs, *RferOR20* and *RferOR22* (Fig. 1). Whether these ORs also detect pheromone compounds remains to be determined, and this knowledge would provide a better understanding of pheromone receptor evolution in Curculionidae. Interestingly, we noticed that the C-terminal parts (amino acids 210-404) of the *RferOR20, RferOR22*, and RferOR1 are highly conserved, suggesting that these ORs have recently evolved. This C-terminal part is predicted to be involved in ligand-binding sites in RferOR1, with noticeably 82% of the binding pocket residues being near the C-terminal end (Table S6), this suggests that *RferOR20* and *RferOR22* may likely have pheromone receptor functions. However, the N-terminal parts (first 100 amino acid residues) are highly divergent; thus, *RferOR20* and *RferOR22* may have evolved different functions that remain to be elucidated. Aggregation pheromones from different weevil species of the subfamily Rhynchophorinae (Table S5) were able to penetrate and interact with the binding pocket of RferOR1 fully, suggesting that the binding pockets of the pheromone receptors from these species are similar. When identified, these receptors could be used alongside RferOR1 to create a robust biosensor array for the early detection of infection in date palm fields. Strikingly, the PRs characterized in the cerambycid beetle, *M. caryae*, ^17^ and the European spruce bark beetle, *I. typographus* ^18^, belong to distantly related OR clades (Fig. 1). This highlights the diversity of pheromone receptors in beetles and weevils and suggests that pheromone receptors appeared at several independent times during Coleoptera evolution.

## Conclusion

With the identification of a pheromone receptor finely tuned to the two components of the RPW aggregation pheromone, our findings represent a significant step forward in understanding the chemosensory mechanisms of the primary enemy of palm trees. Our study also defines the RPW as an essential model for exploring the chemical ecology of palm tree weevils and the evolution of pheromone detection in Coleoptera. By demonstrating that *Drosophila* ORNs can be reliably used for the functional expression of Coleoptera ORs and providing a standardized protocol for OR silencing *via* RNAi, we have opened a new workflow that can be applied to the deorphanization of other insects ORs. Finally, our study provides a characterized OR as a new target for RPW pest control. Our study provides experimental evidence that, on the one hand, silencing *RferOR1* may disrupt palm weevil aggregation in palm tree plantations, ultimately disturbing the reproductive process and decreasing the *R. ferrugineus* population, a promising step for preventing coordinated mass attacks. On the other hand, this OR could also be used for the development of biosensors, an E-nose, and behavior-based robots by exploiting pheromone-RferOR1 interactions for pheromone-based RPW monitoring in the field, allowing the early detection of infestations in date palm fields. The discovery of this OR also opens up the possibility of a ‘reverse chemical ecology’ approach^10,38,39^ based on the screening of new semiochemicals able to interfere with the OR response to the pheromone and ultimately with the behaviors of the RPW in the field.

## Methods

For additional information, see *Supplementary Methods*.

### RferOR Phylogenetic analysis, expression analyses and pheromone pre-exposure experiments

Phylogenetic analysis was performed using previously annotated OR amino acid sequences from *R. ferrugineus* ^13^, *Ips typographus* ^18^, *Megacyllene caryae* ^17^, *Dendroctonus ponderosae* ^40^, *Colaphellus bowringii* (GenBank: ALR72489 - ALR72590) and *Sitophilus oryzae* (GenBank BioProject: PRJNA566109) to identify gene orthologs and paralogs. Multiple sequence alignments were performed using MAFFT v.7 ^41^, with the E-INS-i iterative refinement strategy and default parameters, followed by manual trimming. The JTT+G+F substitution model was determined as the best-fit model of protein evolution based on the AIC by using ProtTest *v*. 3.4 ^42^. Phylogenetic reconstruction and analysis of the OR distribution were performed *via* the maximum likelihood method with 1000 bootstrap replications using RAxML *v*. 8 ^43^.

The expression of 71 *RferORs* previously identified in the antennal transcriptome ^13^ was mapped in the male and female antennae and male snout, legs, thorax, abdomen and wings of 20-day-old adult insects using RT-PCR (*Supplementary Methods*). Expression of the 20 antennae-specific *RferORs* in the male and female antennae of individuals from the laboratory colony and a date palm field was quantified using RT-qPCR (*Supplementary Methods*). Relative expression levels of *RferOR* genes were measured by normalization to that of *tubulin* and *β-actin* ^44^ (Table S1) and calculated using the 2^-ΔΔC^_T_ method. Pheromone pre-exposure experiments were conducted using laboratory-reared RPW adults. Male and female adults were isolated in separate stimulus containers and pre-exposed to a synthetic pheromone blend composed of ferrugineol and ferrugineone (9:1) for 4 h (see details in *Supplementary Methods*). The adults were immediately killed by immersion in liquid nitrogen, and the antennae were dissected. Total RNA was extracted from the antennae, relative expression analysis of the 20 antennal-specific ORs was performed using qPCR and relative expression levels were compared with those in field-collected RPWs (*Supplementary Methods*). Laboratory-reared RPWs that were not exposed to pheromone stimuli were used as a negative control, and changes in OR expression in response to exposure to pheromone were calculated using the 2^-ΔΔC^_T_ method ^45^.

### Gene silencing and RT-qPCR validation

The full-length open reading frame sequences of the six most highly expressed *RferORs* were obtained by amplifying both the 5’ and 3’ cDNA ends using the rapid amplification of cDNA ends technique^44,46^ (*Supplementary Methods*). Full-length double-stranded RNAs (dsRNAs) were synthesized through *in vitro* transcription using the MEGAscript RNAi Kit (Life Technologies, USA) following a previously described method ^44,46,47^ The dsRNA-injected RPW pupae were maintained as previously described ^46^. Nine lines of injected weevils were generated: six lines that were each injected with one of the OR dsRNAs and three control lines. Controls consisted of weevils injected with a universal negative dsRNA control (Integrated DNA Technologies, Leuven, Belgium) (hereafter, negative control), weevils that were not injected (hereafter, NI) and weevils injected with nuclease-free water (hereafter, NFW). The emerging adults were transferred to a separate box containing a piece of fresh sugarcane and maintained for 21 days until their use in behavioral assays and electrophysiology experiments (see details in *Supplementary Methods*). For RT-qPCR verification of transcript knockdown, the antennae of 21-day-old adult weevils were separately dissected, and qPCR was conducted using the methods described above ^44^.

### Olfactometer assay and electroantennography (EAG)

We followed previously described methods^44,46^. For behavioral studies, 15- to 18-day-old adult insects were starved overnight, and the response of each insect to pheromone stimulation was recorded three times (*Supplementary Methods*). For EAG experiments, six 21-day-old adult RPWs were tested per group (dsRNA-injected, NFW-injected and NI). The antennal responses to each stimulus were recorded using a Syntech Acquisition IDAC-2 controller connected to a computer and recorded and processed using EAG 2012 *v*1.2.4 (Syntech, Kirchzarten, Germany) (*Supplementary Methods*).

### Transgenic expression of *RferOR1* in *Drosophila* ORNs

The ORF encoding *RferOR1* was cloned into the *pUAST.attB* vector (*Supplementary Methods*). Transgenic *D. melanogaster UAS-RferOR1* lines were generated by BestGene Inc. (Chino Hills, CA, USA) by injecting the EndoFree *pUAST.attB-RferOR1* plasmid into fly embryos expressing the integrase PhiC31 and carrying an *attP* landing site within region ZH-51C of the second chromosome ^48^. *Drosophila* lines expressing the *RferOR1* transgene in at1 trichoid sensillum ORNs (genotype *w*; *UAS-RferOR1*, *w*^+^; *Or67d*^GAL4^) were generated by crossing the *UAS-RferOR1* line to the *Or67d*^GAL4[2]^ line ^19^. Genomic integration and expression of *RferOR1* in *Drosophila* were verified using PCR and RT-PCR of *Drosophila* genomic DNA and antennal RNA, respectively.

### Single-sensillum recordings and odor simulation

Single-sensillum recordings (SSRs) on the at1 sensilla of 2- to 5-day-old flies were performed following standard procedures ^49^. During SSRs, flies were kept alive under a constant 1.5 L.min^-1^ flush of charcoal-filtered, humidified air delivered to the antenna until the recording finished. A wide range of weevil and palm beetle aggregation pheromone compounds (see details in Table S5) and structurally related chemicals, including the two components of the RPW pheromone (4*RS*,5*RS*)-4-methylnonan-5-ol (>98% purity, ChemTica Int., Costa Rica) and 4(*RS*)-methylnonan-5-one (>98% purity, ChemTica International), were tested to draw the response spectra of ORNs expressing *RferOR1* (*Supplementary Methods*). All molecules were dissolved in *n*-hexane (1 μg/μl, 10 μg deposited on filter paper inside the stimulus cartridge), and cartridges containing only hexane or 10 μg of 11-*cis*-vaccenyl acetate (cVA), the ligand of the *Drosophila* receptor OR67d, were used as controls. For the most potent ligands identified, we performed dose-response analyses with ligands at increasing amounts of 0.001, 0.01, 0.1, 1.0 and 10 μg on filter paper strips. Stimulations lasted 500 ms (see details in *Supplementary Methods*), and the responses of at1 ORNs expressing *RferOR1* were calculated by subtracting the spontaneous firing rate (measured over 500 ms before stimulation) from the firing rate during the stimulation. Responses to the different stimuli were compared to the response to solvent alone using Kruskal-Wallis ANOVA followed by Dunn’s *post hoc* test with Past v.3.26 ^50^.

### Structural modeling and docking of *RferOR1*

The *RferOR1* protein sequence was used to model the 3-dimensional (3D) structure of *RferOR1* using the homology modeling server I-TASSER (https://zhanglab.ccmb.med.umich.edu/I-TASSER/). After that, the Computed Atlas of Surface Topography of proteins (CASTp) web server (http://cast.engr.uic.edu9) was used to identify potential binding pockets in the protein. The CASTp data were analyzed as reported ^37^. EADock DSS software from the SwissDock server provided by the Swiss Institute of Bioinformatics (http://swissdock.vital-it.ch/docking) was used to dock the target ligands into the RferOR1 protein. The resulting docking predictions were viewed and analyzed using the SwissDock server plugin in UCSF Chimera as reported in ^37^ Twenty-nine target ligands, including the two RPW aggregation pheromone components and structurally related chemicals (including those tested on *RferOR1* expressed in *Drosophila* ORNs), were used for the docking experiments (as detailed in Table S8).

### *RferOR1* expression and EAG responses to pheromone according to RPW age

Relative *RferOR1* expression in male and female adult RPWs of different ages (0, 20 and 60 days) was measured using RT-qPCR under the same conditions described above. The antennal responses of male and female adult RPWs of the same age (0, 20 and 60 days) to ferrugineol and ferrugineone were recorded and processed using EAG (Syntech) as described above.

## Acknowledgments

Funding for this research - grant numbers: KACST-NSTIP 12-AGR2854-02 and KAUST-OSR-2018-RPW-3816-1, OSR-2018-RPW-3816-4 of Saudi Arabia. The authors are grateful to the Deanship of Scientific Research, King Saud University, for funding through the Vice Deanship of Scientific Research Chairs. The authors thank Anne-Francoise J. Lamblin and T. A. Abrajano of KAUST-OSR for their invaluable support. The authors thank the date palm farmers in the Al Kharj and Al Qassim areas for their support in obtaining red palm weevils and advice in adult weevil collection.

## Authors’ contributions

BA, EJ.-J, KP, and AP conceived of the study and acquired the grant. JJ, BA, and MAA participated in RPW field collection, rearing and electrophysiology. BA, AP and JJ performed transcriptome, assembly and annotation. BA and JJ performed the silencing experiment. NM, RC, BA, and EJ-J carried out the *Drosophila* functional expression studies. KC, KP, and BA performed the docking experiment. BA wrote the paper with contributions from NM, JJ, EJ-J, KC, and KP.

## Data availability

The *RferOR1* sequence reported in this paper has been deposited in the GenBank database (accession no. MK060009). RferOR2-RferOR20 and RferOR22 sequences are available in the Dataset S1.

## Supplementary Information

This article, containing Supplementary Methods, Supporting Information and Dataset S1.

## Supplementary information

Supplementary Methods

Figures S1 to S5

Tables S1 to S10

Legends for Dataset S1

SI References

**Fig. S1** Tissue-specific expression analysis of 71 ORs identified from *Rhynchophorus ferrugineus*. Tissues used are indicated as AM (male antennae), AF (female antennae), Sn (male snout), Lg (male legs), Thx (male thorax), Ab (male abdomen) and Wg (male wings). Primer details and PCR product sizes are provided in Table S1. *RferOR1* to *RferOR20* - antennal specific ORs.

**Fig. S2.** Highlight of the main binding pocket (active site) of the red palm weevil pheromone receptor (RferOR1) that was identified using CASTp server. In **(a)** and **(b)** the active site is shown within the whole protein and the constituent amino acid residues are labelled in both figures. In (b) the pocket is shown in hydrophobicity surface representation and colored in dark pink. In **(c)** and **(d)** the pocket is shown on its own and the constituent amino acid residues are labelled in both figures. In **(d)** the pocket is shown in hydrophobicity surface representation and colored in dark pink. The figures were rendered using UCSF Chimera software from the potential binding pockets data generated by CASTp server. The characteristic features of the active site are summarized in Table S6.

**Fig. S3.** Mouth openings for the active site of *RferOR1*. **(a)** Mouth openings for the active site of RferOR1, residues making up the entry mouth are shown in magenta, while residues making up the exit mouth are shown in blue, while in **(b)** the mouths are shown interactively (hydrophobicity surface). In RferOR1 the active site channel has a distinct entry mouth and an exit mouth. The residues making up the entry mouth are D77, K78, D298, K301, V302, and D308, while those making up the exit mouth are N174 and S176.

**Fig. S4** Access and interaction of the ligands with the binding pocket (active site) of RferOR1. The ferrugineol is able to penetrate fully into the binding pocket, but VUAA1 cannot. In (**a**) a stick model is illustrated, while in (**b**) the pocket is shown interactively (hydrophobicity surface). The pocket constituent amino acid residues are labelled in **green** in both figures.

**Fig. S5.** Structural analysis, contacts (yellow lines) between ferrugineol ligand and RferOR1 active site residues. In **(a)** the whole pocket is shown, while in **(b)** only residues that have direct contact (van der Waals interaction) with the ligand are shown. The analysis was carried out using the Structural analysis/Find Clashes/Contacts tool of UCSF Chimera.

**Table S1**. List of primers used in tissue expression analysis, gene silencing, *Drosophila* expression and RT-qPCR experiments.

**Table S2.** RPKM values and RT-qPCR based on relative quantification of antennal specific ORs in pheromone pre-exposed (Ind) and field samples **(Fld**). Mean fold changes compared to endogenous control *tubulin* and *β-actin* (5). Expression of antennal specific ORs were quantified and RT-qPCR relative quantification compared to endogenous control. The RPKM values represents data from a pool of antennal samples from male and female insects (2). For relative quantification, 2^-ΔC_T_^ values were calculated for the samples: Pheromone Pre-exposed Male (Ind male), Pre-exposed Female (Ind Female), Field1 Male (Fld1 Male), Field1 Female (Fld2 Female), Field2 Male (Fld2 Male) and Field2 Female (Fld 2 Female) (see *Supplementary Methods*).

**Table S3.** The significance in olfactometer preferences exhibited by each group of insects were analyzed by one-way ANOVA followed by LSD method. P values were compared within groups and provided in the table (alpha level of significance *P*<0.05).

**Table S4**: The significance in EAG responses exhibited by each group of insects were analyzed by Turkey’s HSD method, and P values were compared within groups and provided in the table (alpha level of significance *P*<0.05). Values with significant differences were highlighted in bold.

**Table S5.** List of aggregation pheromone compounds (with the corresponding species) and structurally-related chemicals used to stimulate OSNs expressing RferOR1. All species listed are weevils belonging to the sub-family Rhynchophorinae, except *Oryctes sp*. that are palm tree feeding beetles from the family Scarabaeidae.

**Table S6:** Listing the active site amino acid residues of RferOR1. Residues making up the entry mouth are highlighted **in bold** while those making up the exit mouth are highlighted in bold italic. Residues that identified by PoPMuSiC server to give possibly stable mutants are highlighted in yellow.

**Table S7:** Binding pocket features of RferOR1.

**Table S8:** Listing the compound used for docking experiment.

**Table S9:** Docking Screening results. The actual docking experiments and data analysis were carried out as described in the methods section. *Drosophila* ORNs respond to the following compounds, are highlighted in yellow.

**Table S10:** Identification of interatomic contacts based on VDW (van der Waals) radii between ligand and active site amino acid residues of RferOR1. The analysis was carried out using the Structural analysis/Find Clashes/Contacts tool of UCSF Chimera. #0 = residues in the model that interact with the ligand; while #1.1 = ligand number –there is only one ligand in this case, ferrugineol. ***Contacts*** - all kinds of direct interactions: polar and nonpolar, favourable and unfavourable (including ***clashe***s- unfavourable interactions where atoms are too close together; close contacts). ***Overlap values:*** For detecting contacts, negative cut-off values of 0.0-(−1.0) Å with an allowance of 0.0 Å are generally reasonable (default contact criteria −0.4 and 0.0 Å, respectively). ***Overlap values:*** For detecting clashes, cut-off values of 0.4-1.0 Å and allowance values of 0.2-0.6 Å are generally reasonable (default clash criteria 0.6 and 0.4 Å, respectively).

**Dataset S1:** *RferOR2* to *RferOR20* and *RferOR22* nucleotide sequences.

